# Identification of novel evolutionarily conserved genes and pathways in human and mouse musculoskeletal progenitors

**DOI:** 10.1101/2024.03.14.584954

**Authors:** S Saravanan, A. S Devika, Abida Islam Pranty, R. V Shaji, Raghu Bhushan, James Adjaye, Smita Sudheer

## Abstract

The axial skeletal system and skeletal muscles of the vertebrates arise from somites, the blocks of tissues flanking both sides of the neural tube. The progenitors of Somites, called the Presomitic Mesoderm (PSM) reside at the posterior end of a developing embryo. Most of our understanding about these two early developmental stages comes from the studies on chick and mouse, and in the recent past, there have been a few studies on human. Here, we have analysed and compared the RNA-sequencing data of PSM and somite tissues from Mouse and Human. The functional and pathway enrichment analysis identified the key Hub-genes that are evolutionarily conserved in the PSM and the somites of both the organisms that include 23 multifunctional genes likely to be associated with different developmental disorders in humans. Our analysis revealed that NOTCH, WNT, MAPK, BMP, Calcium, ErbB, cGMP-PKG, RAS and RAP1 signaling pathways are conserved in both human and mouse during the development of PSM and Somites. Furthermore, we validated the expression of representative conserved candidates in the hESCs-derived PSM and somite cells (*NOG*, *BMP2*, *BMP7*, *BMP5*, *HES5* and *MEF2C*). Taken together, our study identifies putative gene interactions and pathways that are conserved across the mouse and human genomes, which may potentially have crucial roles in human PSM and somite development.

## Introduction

Gastrulation initiates with the formation of the primitive node and the primitive streak (PS), which allows the rearrangement of epiblast cells, eventually giving rise to the mesoderm and the endoderm lineages. Presomitic mesoderm (PSM), the progenitors of somites that give rise to the axial skeletal system and skeletal muscles, originates in the PS and resides in the posterior end of a developing embryo ^1^. The expression of the T-Box Transcription factors, Brachyury (*T*) ^2^, *Tbx6*^3^ and Mesogenin 1 (*Msgn1*) ^4,5^ and the oscillation of clock genes involved in the segmentation clock are the hallmarks of PSM ^4,6^. According to the regulated activity of two independent gene regulatory networks, known as the segmentation clock and the wavefront phenomenon, the mesenchymal PSM cells form new pairs of somites in the anterior end of the PSM ^7–9^. The differentiation occurs from the anterior to posterior direction, while the migration of progenitor cells occurs in the posterior to anterior direction. During this process, the pre-segmented PSM remains in the caudal region, and the segmented PSM resides in the rostral region, dividing the PSM into the posterior PSM and the anterior PSM respectively.

Somites pinch off from the rostral end of the PSM in response to the signaling pathways involved in the clock and wavefront phenomenon. The mutually antagonistic activity of the FGF and the retinoic acid (RA) signaling gradients involved in the wavefront model creates a zone of determination where the mesenchymal PSM cells become compacted to form somitomeres ^10–12^. These somitomeres undergo mesenchymal to epithelial transition to form somites, with an outer epithelial layer and an inner mesenchymal core. The exposure of various signaling pathways from the surrounding cells induces the differentiation of the nascent somites into the ventromedial sclerotome and the dorsolateral dermomyotome ^13,14^. The *Pax1* and *Pax9* positive sclerotome differentiates into the vertebral column and intervertebral disc, where the *Pax3* and *Pax7* positive dermomyotome develops into the skeletal muscle and dermis.

Here, we have analysed the *in vivo*-derived whole transcriptome data of human and mouse PSM and Somites and identified the evolutionarily conserved known and putative signalling pathways and hub genes, that could potentially have crucial roles in the development of these cell types and their descendants.

## Results

### Putative regulatory hub genes of mouse interact with the PSM markers, T and MSGN1 and are involved in the signalling pathways, Rap1, PI3K-AKT, MAPK, Hippo, RAS, WNT

Publicly available data of PSM and somites from the mouse E8.25 embryonic stage embryos (E-MTAB-6155) ^15^ was utilized to find out the hub-genes involved in the PSM and somites. Principal Component Analysis (PCA) of the differentially regulated genes shows that, the posterior PSM (PSM1, PSM2 and PSM3) were clustered together and segregated from the anterior PSM (PSM4 and (PSM5) and somites (Figure 1. a). The most posterior part of the PSM (PSM1) and the somites are the most variant in this trajectory (Figure 1. a).

**Figure 1.**
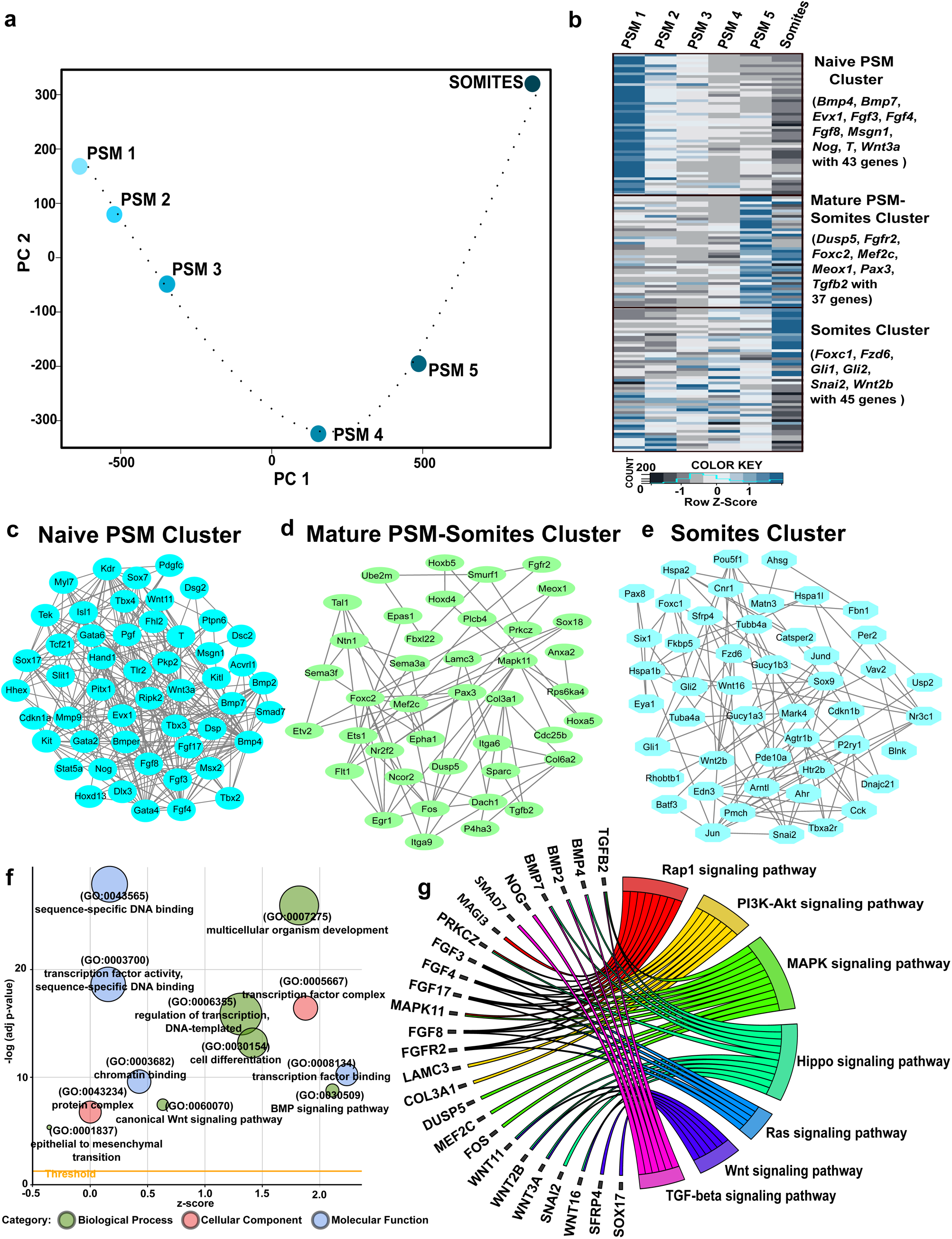
Gene expression and regulation in the mouse PSM and somites (a) PCA of mouse PSM from posterior to anterior axis (PSM 1 to PSM5) and somites. Hub-genes were identified from the gene regulatory network analysis of the selected WGCNA clusters (b) Heatmap showing the expression status of Hub-genes identified from the selected clusters (c) Naïve PSM cluster (Black (Figure S1)) (d) Mature PSM-Somites cluster (Red (Figure S1)) and (e) Somites cluster (Yellow (Figure S1)). Functional enrichment analysis of the Hub-genes shows their involvement in different (f) biological, cellular and molecular functions and in (g) signaling pathways

We performed weighted correlation network analysis (WGCNA) with the differentially expressed genes (DEGs) and selected three clusters (Black cluster, Red cluster and Yellow cluster), based on the presence of known markers of PSM and Somites in these clusters. Gene regulatory networks (GRN) were constructed using STRING, visualized using Cytoscape and hub-genes were identified (Figure 1. b – d, Table S1) ^16–18^. Hub-genes are the genes with high connectivity or correlation in a module. Based on the expression of the hub genes, the selected clusters were identified to represent the most posterior end of the PSM (PSM1) (black cluster: the naïve PSM cluster), the anterior-most part of the PSM (PSM5) (red cluster: the mature PSM-Somite cluster) and the Somites (yellow cluster: the mature Somite cluster) (Figure 1. b-e, Table S1). Chip-seq data available from the previously reported studies clearly shows that the candidates in the identified Hub-genes interact with the PS and the PSM markers, T and MSGN1 ^4,19^ which validates our predictions. Based on this, the pan-mesoderm marker, *T* ^19^ interacts with some of the Naïve PSM cluster genes (*Bmp4*, *Fbln2*, *Fgf17*, *Fgf8*, *Hhex*, *Msx2* and *Wnt3a*), the Mature PSM-somite cluster genes (*Cdc25b*, *Epas1*, *Epha1*, *Meox1*, *Prkcz* and *Sox18*) and the mature Somite cluster genes (*Cck* and *Foxc1*) (Figure 1. c-e, Table S1). The PSM marker, *MSGN1* ^4^ interacts with several genes from the three identified Hub-gene clusters: *Fbln2*, *Gata4*, *Myl7*, *Tbx3*, *Gata6* and *Slit1* (Naïve PSM cluster), *Atp8a1*, *Epas1*, *Sparc*, *Flt1*, *Dach1*, *Epha1*, *Fgfr2*, *Msi1*, *Pax3*, *Plcb4* and *Rhof* (Mature PSM-Somite cluster), *Eya1*, *Foxc1*, *Myl1*, *Rhobtb1*, *Six1*, *Ahsg*, *Blnk*, *Fkbp5*, *Gucy1a3*, *Magi3*, *Tbxa2r*, *Tubb4a*, *Wnt2b* (Somite cluster) (Figure 1. c-e, Table S1).^3,17^.

The functional enrichment analysis of the identified Hub-genes shows that, these genes are important for species-specific DNA-binding, multicellular organism development, transcription factor activity, transcription factor complex activity, cell differentiation, etc. (Figure 1. f, Table S1). The pathway enrichment analysis indicates the involvement of the Hub-genes in various signaling pathways such as Rap1, PI3K-AKT, MAPK, Hippo, RAS, WNT (Figure 1. g, Table S1), which are important for the development and further differentiation of PSM.

### T, SALL4 and LEF1 among the hub genes interacting with the PSM markers, TBX6 and MSGN1 and the conservation of Calcium signalling in human somite development

The whole transcriptome dataset of human PSM, somites and developed somites from human embryos of age 4.5–5 weeks of gestation (GSE90876) ^20^ was used for the identification of DEGs and Hub-genes involved in the development of human musculoskeletal progenitors. The cluster dendrogram indicates the developmental progression of musculoskeletal progenitors from PSM to somites and further into developed Somites (Figure 2. a).

**Figure 2.**
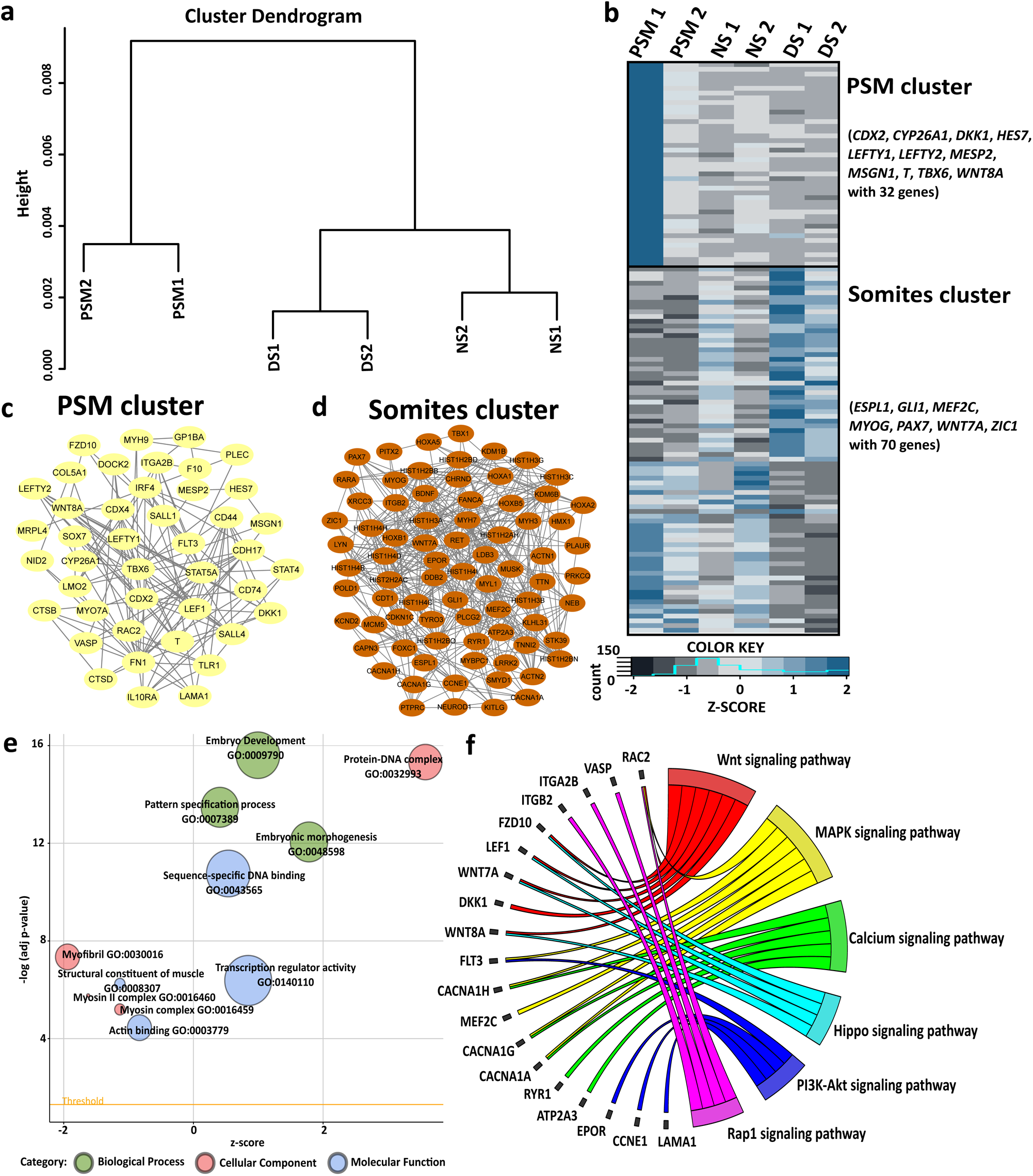
Gene expression and regulation in the human PSM and somites (a) Cluster dendrogram of human PSM, somites and developed somites. From the Gene regulatory network analysis of selected WGCNA clusters, Hub-genes were identified (b) Heatmap indicating the expression status of Hub-genes identified from the selected clusters (c) PSM cluster (Yellow (Figure S2)) (d) Somite cluster (Brown (Figure S2)). Functional enrichment analysis of the Hub-genes shows their involvement in different (e) biological, cellular and molecular functions and in (f) signaling pathways

DEGs were subjected to WGCNA clustering, and two clusters were selected (Yellow cluster and Brown cluster), in which the known markers of PSM and somites were clustered (Figure S2). The members of yellow and brown clusters represent the upregulated genes in PSM and in somites respectively (Figure 2. b – d, Table S2). The Hub-genes in the yellow cluster (PSM cluster) contains the PSM markers *TBX6* and *MSGN1* and important genes such as *T*, *MESP2*, *CYP26A1*, *HES7*, *WNT8A*, *SALL4*, *LEF1*, etc. which are expressed or involved in the development of mesoderm or PSM (Figure 2. c, Table S2) ^3,5,21–24^. In mouse, SALL4 is important for the maintenance of neuromesodermal progenitors and the proper development of PSM cells ^25^. The SALL4 knockout negatively effects the expression of PSM associated genes *T*, *Lef1*, *Msgn1* and *Hes7* ^25^, and our analysis shows its probable conservation in human somitogenesis. The Hub-genes identified from the brown cluster (Somite cluster) contains somite-associated genes such as *FOXC1*, *MEF2C*, *MYOG*, *PAX7*, etc. (References) (Figure 2. d, Table S2).

The functional enrichment analysis shows that the PSM and Somite Hub-gene clusters are involved in embryo development, transcription regulator activity, embryonic morphogenesis, species-specific DNA-binding, protein-DNA complex, structural constituent of muscles, etc. (Figure 2. e, Table S2). The pathway enrichment analysis indicates the role of the predicted Hub-genes in various signaling pathways such as WNT, MAPK, Calcium, Hippo, PI3K-AKT and Rap1 which have crucial roles in musculoskeletal progenitor development (Figure 2. f, Table S2). The importance of Calcium signalling in somitogenesis has been deciphered in Zebra fish ^26,27^. Calcium signalling is downstream of FGF signalling ^28,29^, having a prime role in PSM development ^26,27,30^. Till date, there is no direct evidence of the involvement of Ca signalling in mouse and human, however, the appearance of Ca signalling genes among the hub genes shows its possible role in human and mouse somitogenesis.

### Muscle development genes and the signalling pathways, NOTCH, WNT, MAPK, Calcium, ErbB, cGMP-PKG, RAS and RAP1 are evolutionarily conserved in somitogenesis of mouse and human

To identify the evolutionarily conserved genes involved in mouse and human musculoskeletal progenitor development, the differentially expressed genes (DEGs) between mouse and human were compared (Figure 3. a). A total of 1670 genes were commonly regulated in both the organisms (Figure 3. a, Table S3). Further, the functional enrichment analysis of these 1670 genes in human and mouse databases reveals that the genes are involved in various biological processes such as, Striated muscle tissue development (*DKK1*, *BMP2*, *NOG*, *KLF4*, *BMP7*, *BMP5, T*, *Bmp7*, *Dll1*, *Nog*, *Bmp5*, *Mef2c*), skeletal muscle tissue development (*DLL1*, *DKK1*, *KLF5, Mef2c*, *Dkk1*, *Dll1*), muscle tissue development (*DKK1*, *BMP2*, *NOG*, *KLF4*, *BMP7*, *BMP5, T*, *Bmp7*, *Dll1*, *Nog*, *Bmp5*, *Mef2c*), muscle organ development (*DKK1*, *BMP2*, *TCF15, Mef2c*, *Tcf15*, *Dkk1*, *Nog*), embryonic organ development (*PAX8*, *MEF2C, Cdx4*, *Cdx2*, *Nog*, *Bmp5*, *Dll1*, *Pax8*, *Zic3*) and anterior/posterior pattern specification (*MESP2*, *CDX4*, *MSGN1*, *TBX6*, *CDX2*, *BMP2*, *HES5, Cdx4*, *Cdx2*, *Msgn1*, *Tbx6*, *T*, *Dkk1*, *Zic3*, *Meox1*, *Tcf15*, *Bmp2*, *Hes7*, *Nog*) (Figure 3. a, Table S4). By analysing the list of genes under various biological processes in mouse and human, we found that most of these genes have important roles in the induction of PSM, somitogenesis and the maturation of somites ^31–34^.

**Figure 3.**
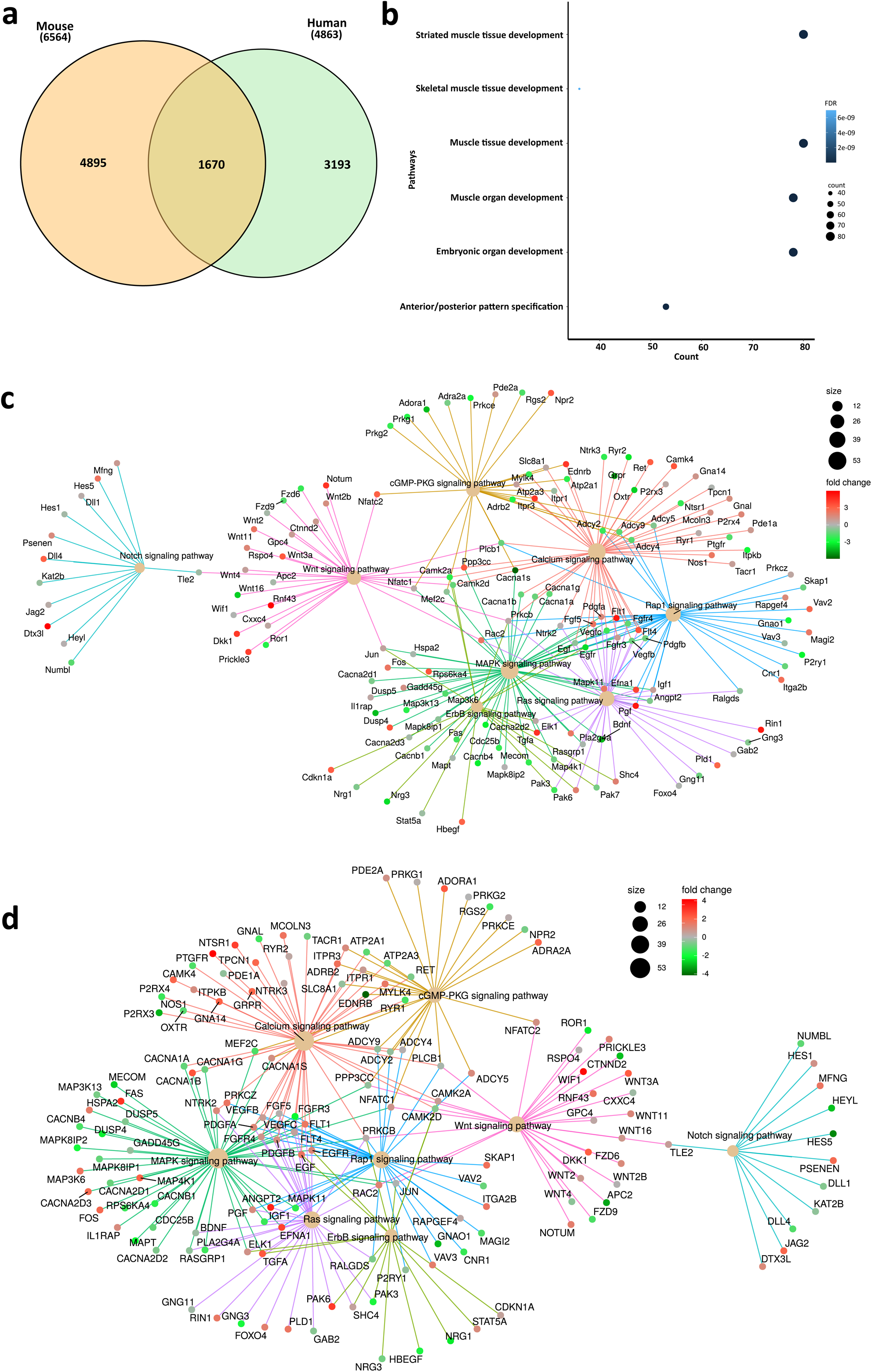
Evolutionarily conserved gene and pathway interaction network (a) Venn diagram of mouse and human DEGs, (b) Biological process for evolutionarily conserved genes in human, (c) Pathway interaction network of evolutionarily conserved genes in mouse, (d) Pathway interaction network of evolutionarily conserved genes in human

Pathway enrichment analysis was carried out for the commonly regulated 1670 genes and pathway interaction network was constructed with the most significant signaling pathways that were common for both the organisms (Figure 3. c-d, Table S4). The commonly regulated pathways include NOTCH, WNT, MAPK, Calcium, ErbB, cGMP-PKG, RAS and RAP1 signaling. In a developing embryo, FGF, WNT and NOTCH signaling pathways (Figure 3. c-d) interact with *T*, *Tbx6* and *Msgn1* and promotes the differentiation and maintenance of musculoskeletal progenitor ^5,35–38^. RAS-MAPK/ERK1/2 signaling cascade is an effector of FGF pathway, important for early embryonic development ^39,40^. FGF signaling is involved in somitogenesis and is highly active in the posterior side of a developing embryo which maintains a crosstalk with WNT and NOTCH signaling pathways to sustain the progenitor population in the tail bud ^41–45^. Transcriptome data of known and putative clock genes involved in somitogenesis shows that the genes involved in MAPK (*PDGFA*, *NFATC1*, *TGFA*, *DUSP4* and *EFNA1*), RAS (*EFNA1*, *PDGFA*, *TGFA*, *BDNF* and *Foxo4*), RAP-1 (*EFNA1*, *VAV2* and *PDGFA*), WNT (*NFATC1*, *WNT11*, *DKK1*), NOTCH (*Hes5*, *Hes1*, *Dll1*) signaling pathways oscillate during somitogenesis (Figure 3. c-d, Table S4) ^32^. In addition to this, calcium, ErbB and cGMP-PKG signaling pathways are important for gastrulation in embryos, differentiation of mesodermal lineages and somitogenesis (Figure 3. c-d, Table S4) ^26,27,46–49^. Taken together, from this analysis, we have identified putative genes, pathways, and pathway interaction networks, which are probably conserved among mouse and human skeletal progenitors.

### Evolutionarily conserved multifunctional genes involved in the musculoskeletal progenitor development

Evolutionarily conserved multifunctional genes were predicted based on the functional and pathway enrichment analysis of the common 1670 genes identified from the DEGs of mice and humans (Figure 4. e-f, Table S5).

**Figure 4.**
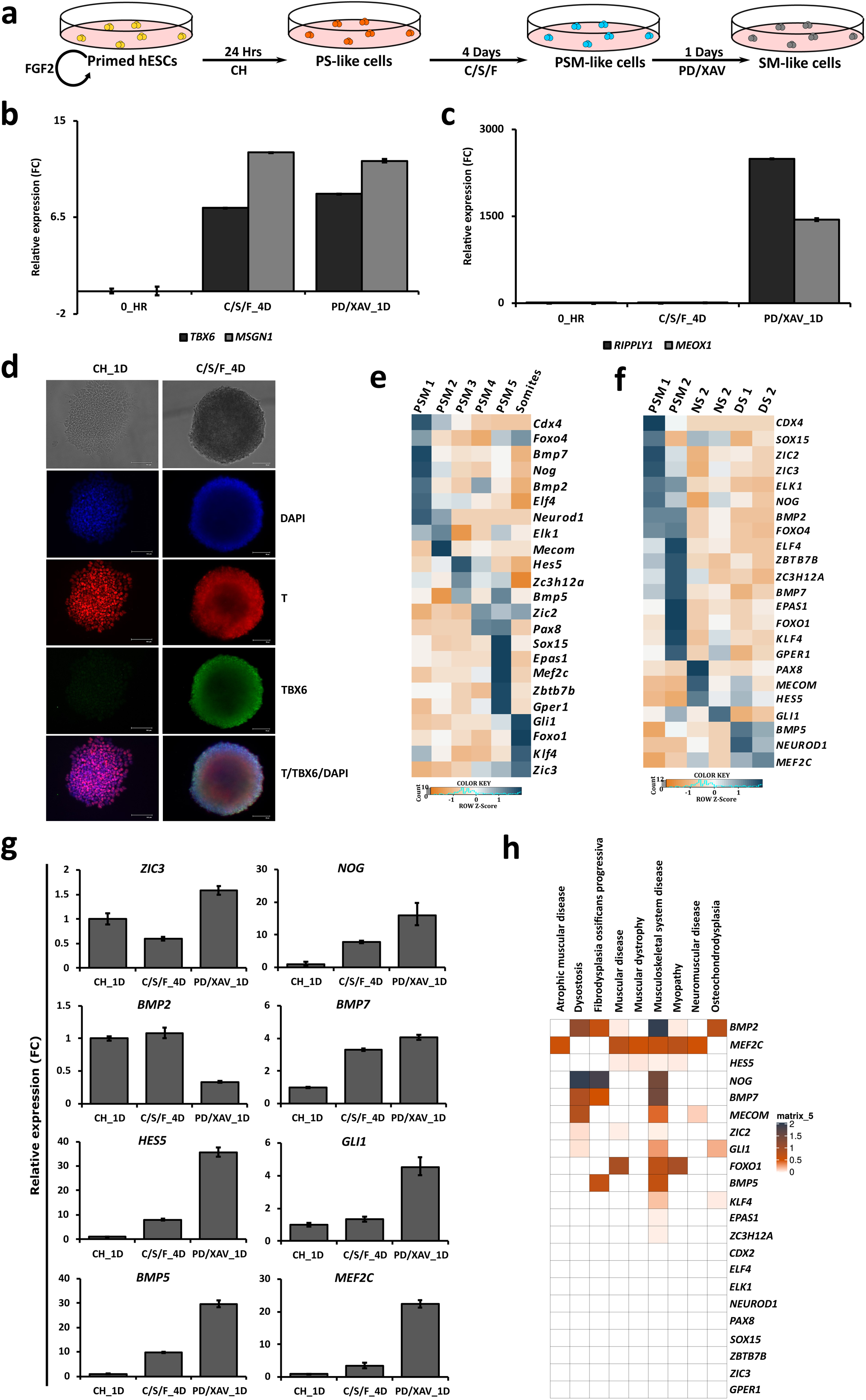
Evolutionarily conserved multifunctional genes and its validation using *in vitro* derived musculoskeletal progenitors from hESCs (a) schematic representation of *in vitro* derived musculoskeletal progenitors from hESCs (b-c) RT-qPCR of PSM and somite markers of indicated samples. Data are Mean ± s.d., n=2, (d) Immunocytochemistry image of T and TBX6 in indicated samples, (e-f) Heatmap showing the expression status of the predicted evolutionarily conserved multifunctional genes in (e) mouse and (f) human (g) RT-qPCR validation of the predicted evolutionarily conserved multifunctional genes in *in vitro* derived human musculoskeletal progenitors. Data are Mean ± s.d., n=2, (h) Heatmap represents the involvement of multifunctional genes in indicated developmental impairments based on their corresponding evident score. Graphs shown in (b-c) and (g) are representatives of two independent technical replicates.

Multifunctional genes are the genes that are associated with more than one function and/or signaling pathway. Such genes tend to be more conserved and associated with human disorders^50^. The identification of multifunctional genes can help us better understand the molecular and functional organization of a cell type. From the gene enrichment analysis of the commonly regulated evolutionarily conserved 1670 genes, 23 multifunctional genes were identified (Figure 4. e-f, Table S5).

To validate the expression of the identified 23 multifunctional genes, human Pluripotent Stem Cells (hPSCs) were differentiated into musculoskeletal progenitors (PS, PSM and somites) (Figure 4. a). The hESCs-induced PS (*EOMES* and *T*: Figure S4 A), PSM (*T*, *TBX6* and *MSGN1*: Figure 4. b, d, Figure S4 A) and Somites (*MEOX1*, *MESP2*, *RIPPLY1* and *DLL1*: Figure 4. c, Figure S4. B) were marked and validated by the expression of their representative markers.

From the identified multifunctional genes, the expressions of 8 genes (*ZIC3*, *NOG*, *BMP2*, *BMP7*, *HES5*, *GLI1*, *BMP5* and *MEF2C*) were validated in the hPSC-derived PSM and somite cells (Figure 4. e-g, Table S5). *Hes5*, *Zic3*, *Zic2* and *Foxo4* are important for mesoderm and neural differentiation (Figure 4. e-g, Table S5) ^51–54^. *HES5*, *ZIC3* and *Zic2* have a crucial role in the migration of PS cells during gastrulation and in the segmentation clock, the gene regulatory network involved in somitogenesis (Figure 4. e-g, Table S5) ^32,52–54^. MEF2C, a member of the MEF2 transcription factor family which regulates several skeletal muscle-specific genes is an early marker for somitogenesis (Figure 4. e-g, Table S5) ^55^. *GLI1* (Figure 4. e-g, Table S5), an intracellular signaling transducer, and a transcriptional effector of the Sonic hedgehog (Shh) signaling pathway, is expressed in the neural tube and paraxial mesoderm in the developing embryo (Figure 4. e-g, Table S5) ^56^. The BMP and FGF signaling pathways act antagonistic to each other in the developing embryo during PS formation and BMP signaling is important for somite maturation ^57^. The members of the BMP signalling, *NOG*, *BMP2*, *BMP7* and *BMP5* were part of the multifunctional genes, and they were also expressed in the hPSC-derived musculoskeletal progenitors (PS, PSM and somites) (Figure 4. e-g, Table S5). The dysregulation of several of the identified multifunctional genes such as *GLI1, HES5, NOG*, *BMP2*, *BMP7*, *BMP5, MEF2C* have implications in the human musculoskeletal developmental disorders, muscular dystrophy, Osteochondrodysplasias and Spondylocostal dysplasia (Figure 4. h, Table S5). Taken together, we have identified evolutionarily conserved multifunctional genes that are regulated during the development of human musculoskeletal progenitors (PS, PSM and somites), with crucial roles in development, having possible implications for human skeletal developmental disorders.

## Discussion

In post-implantation embryos, the crosstalk between several gene regulatory networks promotes the differentiation of PSM into somites, the progenitors of the musculoskeletal tissue. The majority of our understanding about PSM and somites comes from the studies on Zebra fish, chick and mouse and there have been only a few studies on human. Therefore, we set out to identify the evolutionarily conserved genes in human PSM and Somites, by analysing and comparing the whole transcriptome data from these two tissues of human with mouse (E 8.25) and human (4.5– 5 weeks of gestation) ^15,20^. Here, we identified known and putative genes (hub-genes), signalling pathways and interactions in the human musculoskeletal progenitors, PSM and somites. Finally, our analysis led to the identification of evolutionarily conserved multifunctional genes which have been reported to have implications in human skeletal muscle developmental disorders, such as muscular dystrophy, Osteochondrodysplasias and Spondylocostal dysplasia (Table S5).

In the developing blastula, the crosstalk between BMP, WNT and NODAL signaling creates an anterior–posterior gradient of NODAL and WNT signaling in the epiblast which results in the localized expression of the pan-mesodermal marker *T* and the formation of PS ^43,58–61^. The high concentration of Wnt3a in the PS and the tail bud regulates the expression of Fgf8 and promotes epithelial mesenchymal transition (EMT) in the progenitor cells ^62,63^. FGF pathway components *Fgf3*, *Fgf4*, *Fgf8* and *Fgf17* are expressed in the tail bud, the mRNA of Fgf3, Fgf8 and Fgf17 creates posterior to anterior gradient with in the PSM and Fgf4 is localized in anterior part of the PSM ^45,64–66^. The progenitor cells in the tail bud undergoes EMT and moves from the posterior to anterior end of the embryo according to the gradient created by FGF8 and FGF4 thus helps in the axial elongation in embryos ^67,68^. In developing embryo, WNT and FGF signaling pathway acts antagonistic to retinoic acid (RA) pathway by inducing the expression of retinoic acid-metabolizing (inactivating) enzyme CYP26a1 in the posterior PSM ^45,69,70^.

WNT and FGF signaling pathways promotes the expression of paraxial mesoderm specific genes, *T*, *Tbx6* and *Msgn1* ^71–73^. The elevated level of T in PSM cells regulates the expression of Wnt3a and Cyp26a1 in the posterior end ^74^. WNT and FGF signaling pathway together with CYP26a1 limits the concentration of RA in the posterior end and creates an anterior to posterior gradient of RA signaling in developing embryo and helps in the migration of PSM cell towards anterior end ^45,69,70,75,76^. The reduced activity of FGF pathway from posterior to anterior end created by the opposing activity of RA signaling pathway negatively affects the expression of *Snai* genes in anterior PSM and promotes the expression of integrins or cadherins ^42,77,78^. During somitogenesis, Wnt3a act upstream of *Fgf8*, *Dll1*, and *Ctnnb1* components of FGF, NOTCH and WNT signaling pathways respectively and also promotes the expression of negative regulators of the WNT signaling pathway, *Axin2* and *Dkk1* ^41^. FGF and NOTCH signaling maintain their oscillations during somitogenesis by promotes the expression of ERK inhibitors *Dusp4*, *Dusp6*, and *Spry2* and the transcriptional repressor, *Hes7* ^6,79–83^. The oscillatory activity of the genes involved in these signaling pathways together creates a zone which promoter the formation the new pair of somites called the determination front characterized by the expression of (Mesp2, Pax3, Foxc1/2, and Meox1/2) ^43^. The PSM marker TBX6 together with NOTCH signaling pathway promotes the expression of *Mesp2* in the anterior side of the determination front and creates a positive and negative regulatory loop between TBX6, MESP2 and the NOTCH ligand, Dll1 ^10,84–87^. *Pax3*, the marker of segmented mesoderm is regulated by the transcription factors MESP2 and PARAXIS expressed in anterior PSM ^88^. The WNT ligands WNT3A, WNT7A and WNT8C secreted from the neighbouring tissues also promotes the expression of Pax3 and Pax7 in developing dermomyotome ^89,90^.

A comparative approach was used to identify the common DEGs involved in the development of paraxial mesoderm in mouse and human. The gene enrichment analysis indicates that the identified genes were involved in skeletal muscle development and in the signaling pathways involved in this process. The pathway interaction network constricted with the evolutionarily conserved genes includes NOTCH, WNT, MAPK, Calcium, ErbB, cGMP-PKG, RAS and RAP1 Signaling pathways shows the crosstalk between these genes during musculoskeletal development. FGF and WNT signaling pathway collectively regulates the convergent extension (CE) or the cell movement during gastrulation. CE helps in the body axis elongation and the morphogenesis in developing embryos ^91^. ErbB signaling pathway is an upstream regulator of PI3K and FGF signaling pathway and its effector, RAS-MAPK/ERK1/2 signaling pathway ^49,92^. In gastrulating embryo, ErbB signaling pathway regulates CE through MAPK and PI3K signaling pathway ^49^. RAS-MAPK/ERK1/2 and protein kinase C (PKC)/Ca^2+^ signaling pathways, the downstream effectors of FGF signaling pathway. Sprouty and Spred, proteins which modulates the protein kinase C (PKC)/Ca^2+^ and RAS-MAPK/ERK1/2 signaling pathways respectively ^28,29,93^. During early gastrulation, Sprouty inhibits protein kinase C (PKC)/Ca^2+^ signaling pathway, hence the RAS-MAPK/ERK1/2 signaling pathway will be active and helps in cell movement in mesoderm formation ^28,29,93^. Alternatively, Spred inhibits RAS-MAPK/ERK1/2 signaling pathway during mid to late gastrulation and turns the activity of FGF pathway through protein kinase C (PKC)/Ca^2+^, which helps in the morphogenesis ^28,93^.

The evolutionarily conserved multifunctional genes identified to be conserved in both mouse and human are involved in several biological, cellular, and molecular functions. Most of them are known to be involved in the development of paraxial mesoderm and in the regulation or regeneration of skeletal muscles and its progenitors ^94–99^. The dysregulation of these genes may cause developmental or functional impairments of musculoskeletal system. NOTCH and BMP/NODAL/ACTIVIN/TGFβ signaling pathways have crucial roles in the formation, maintenance, and differentiation of musculoskeletal and neuronal progenitors. The abnormalities in these signaling pathways lead to several musculoskeletal and neuromuscular impairments such as osteochondrodysplasia, spondylocostal dysplasia, spinal and bulbar muscular atrophy, etc. ^52,100–102^. The identified multifunctional genes such as *MEF2C*, *MECOM*, *ZIC2*, *GLI1*, *FOXO1*, *KLF4* are involved in the normal development and differentiation of musculoskeletal progenitors and their developmental impairments by interacting with various signaling pathways or involved in the transcription of lineage specific markers ^103–116^. Taken together, the Hub-genes and multifunctional genes identified from the musculoskeletal progenitors of mouse and human are involved in the development and differentiation of paraxial mesoderm. Among the multifunctional genes, we have identified 23 genes conserved between human and mouse, that are crucial for embryonic development, interact with several signaling pathways and when dysregulated, lead to skeletal developmental disorders.

## Materials and Methods

### Data collection for meta-analysis

The whole transcriptome data of PSM and somites from the mouse embryos (E 8.25) were obtained from the ArrayExpress database ^117,118^. The gene expression of posterior to anterior PSM and somites from four different mouse embryos (E-MTAB-6155) (https://www.ebi.ac.uk/arrayexpress/experiments/E-MTAB-6155/samples/?query=presomitic+mesoderm or https://www.ebi.ac.uk/biostudies/arrayexpress/studies/E-MTAB-6155) were considered for this study (Ibarra-Soria et al., 2018). In mouse, the RNA sequencing data of PSM were obtained from the five individual segments from the left and right sides of posterior to anterior axis within the tail bud region ^15^(Ibarra-Soria et al., 2018). Gene expression data of human PSM and somites were obtained from Gene Expression Omnibus (GEO) ^119^. The human RNA sequencing data were obtained from PSM, somites and developed somites from two different human embryos of age 4.5–5 weeks of gestation (GSE90876) (https://www.ncbi.nlm.nih.gov/geo/query/acc.cgi?acc=GSE90876) ^20^. The Human and Mouse whole genome and Gene transfer file (GTF) files were collected from the NCBI database.

### Quality Check and Mapping of RNA seq data

FASTQC Version 0.11.5 was used to find out the GC content, total sequence length, and the base sequence quality of each sample. Unlike the original article ^20^, for indexing the human genome we used HISAT2 (Version 2.1.0) ^120^. The Mapping of the indexed Mouse and Human genome was also carried out by HISAT2. Cufflinks (Version 2.2.1) ^121^ used to assembles the transcripts for the RNA-seq samples, where we have found out the FPKM (Fragments per Kilobase of exon per million mapped fragments) for each sample.

### Principle component analysis (PCA) and hierarchical clustering

Principle component analysis (PCA) was done using R package, in which the similarities and dissimilarities between the samples and its replicates were plotted. Using FPKM values, the hierarchical clustering analysis was conducted to show the similar gene expression status of the samples using R packages.

### Differential expression analysis

To find out the differently expressed genes involved in musculoskeletal progenitor development, we used DESeq (Version 1.26.0). Using the raw read counts, the gene expression between samples were identified and filtered based on P-value < 0.05 and the upregulated and downregulated genes were filtered with a threshold of the log2 fold change ≥1 and ≤-1 respectively.

### Co-expression Network Construction

Using DEGs identified from the human and mouse data sets, weighted gene co-expression network analysis (WGCNA) with R packages (Version 1.70.3) was performed to find out the modules cluster of genes that are highly correlated. The modules with genes clustered along with the known markers of musculoskeletal progenitors were considered for the further analysis.

### Identification of Hub-genes and Functional Annotation

Hub-genes are the genes which shows high connectivity or correlation between the genes in the candidate module. To identify the hub-genes, a protein-protein interaction network (PPI) was constricted with search tool for the retrieval of interacting genes (STRING) ^16,122^ and visualized using Cytoscape (Version 3.8.2) ^17^. The hub-genes used for the PPI were selected based on “OR” condition on Betweenness >10, closeness <0.001, and Degree >2. The functional enrichment analysis was performed using the Database for Annotation, Visualization, and Integrated Discovery (DAVID) ^123^.

### Heat map and plot construction

The expression status of each gene in each sample were represented in heatmap constricted using Gplot (R package, Version 3.1.3). The circus plot representing the involvement of hub-genes in various signaling pathways and the bubble plot were generated with GOplot (R package, Version 1.0.2). The bubble plot representing the functional enrichment analysis was constructed against the genes counts and the p-values of each identified function.

### Pathway interaction network construction

Functional and pathway enrichment analysis were performed for evolutionarily conserved genes identified from mouse and human using ClusterProfiler (R package, Version 3.14.3) (https://bioconductor.org/packages/release/bioc/html/clusterProfiler.html). The representatives of functional enrichment analysis were visualized using dotplot. The results obtained from pathway enrichment analysis was represented as pathway interaction network using cnetplot.

### Maintenance and Differentiation of human pluripotent stem cells (hPSCs)

The human embryonic stem cell (hESCs) (BJNhem19 (JNCASRe001-A)) line was procured from Jawaharlal Nehru Centre for Advanced Scientific Research (JNCASR), India. The human-induced pluripotent stem cell (hiPSCs) (D14C2) line was a kind gift from Dr. R. V. Shaji, Centre for Stem Cell Research, (CSCR), InStem, India. hESCs and hiPSC was maintained on vitronectin (VTN) (Gibco, A14700) in presence of Essential 8™ (E8) Medium (Gibco, A1517001). The cells were routinely passaged in 1:6 ratio in every 4 days using 0.5 mM EDTA (Gibco, 15575020) solution during maintenance.

For PS induction, hESCs were exposed to CHIR99021 (CH), inhibitor of GSK-3β for 24 hours and marked by the expression of *EOMES* and *T* (Figure 4. a, Figure S4 A). After PS induction, the cells were exposed to CH (GSK-3β inhibitor), SB431542 (SB) (ALK 4/5/7 inhibitor) and bFGF (C/S/F) for 4 days and detected the expression of PSM markers *TBX6* and *MSGN1* together with the pan-mesodermal marker T (Figure 4. a-b, Figure S4 A). Due to C/S/F treatment for 4 days, the expression the endoderm marker *EOMES* were downregulated, and the expression of pan-mesodermal marker *T* remains unaffected (Figure S4. A). PSM was further differentiated to somites using FGFR inhibitor PD173074 (PD) and WNT pathway inhibitor XAV939 (XAV) ^32^ and confirmed by the expression of *MEOX1*, *MESP2*, *RIPPLY1* and TCF15/PARAXIS (Figure 4. a, c, Figure S4. B).

### Real-Time PCR analysis

Total RNA was isolated using QIAzol Lysis Reagent (QIAGEN, 79306) according to manufactures’ instruction, followed by quantification using NanoDrop Spectrophotometer (Thermo Fisher Scientific). Reverse transcription was performed with the iScriptTM cDNA Synthesis Kit (Bio-Rad, 1708891). Quantitative Real-Time PCR (qRT-PCR) was done using PowerUp™ SYBR™ Green Master Mix (2X) (Applied Biosystems, A25776) with gene-specific primers (Table S6) in a thermal cycler (Roche Light Cycler 480). Data was analysed using the ddCt method, with the house-keeping gene, *ACTB*.

## Data availability

All data utilized for this study are publicly available data sets from previous publications (E-MTAB-6155, GSE90876)^15,20^. The data that supports the findings of this study are available within this manuscript and in supplementary documents.

## List of Supplementary tables

**Table S1:** Hub-genes; List of Hub-genes identified as Naïve PSM cluster, Mature PSM-Somite cluster and the Somite cluster – Include the details of GRN, pathway and functional enrichment analysis

**Table S2:** Hub-genes; List of Hub-genes identified as Naïve PSM cluster, Mature PSM-Somite cluster and the Somite cluster – Include the details of GRN, pathway and functional enrichment analysis

**Table S3:** List of commonly regulated genes in Mouse and Human

**Table S4:** Pathway and functional enrichment analysis of commonly regulated genes in Mouse and Human

**Table S5:** List of Evolutionarily conserved multifunctional genes and their involvement in developmental impairments

**Table S6:** List of Primers

## Acknowledgements

This work was supported by the Department of Science and Technology/ Science and Engineering Research Board (DST/SERB) (ECR/2017/001216), Indian Council of Medical Research (ICMR) (2019-3008/SCR/ADHOC/BMS) and the Council of Scientific & Industrial Research (CSIR) (09/1108(13739)/2022-EMR-1), New Delhi, India.

J.A. acknowledges the medical faculty of Heinrich Heine University for financial support.

## Authors’ contributions

SA.S, D.A.S. designed the study with SM.S, performed the experiments, analysed, and interpreted data. SA.S processed the RNA sequencing data. S.R.V gifted the iPSCs. SA.S and D.A.S. wrote the manuscript and composed the figures. A.I.P, R.B, J.A and SM.S reviewed and edited the manuscript. SM.S conceptualised and supervised the work, acquired funding. SM.S and J.A: final approval of the manuscript.

## Competing interests

The authors declare that they have no competing interests.

## Notes

### Competing Interest Statement

The authors have declared no competing interest.

## References

1 Bénazéraf B, Pourquié O. Formation and segmentation of the vertebrate body axis. Annu Rev Cell Dev Biol 2013; 29: 1–26.

2 Herrmann BG, Labeit S, Poustka A, King TR, Lehrach H. Cloning of the T gene required in mesoderm formation in the mouse. Nature 1990; 343: 617–622.

3 Chapman DL, Agulnik I, Hancock S, Silver LM, Papaioannou VE. Tbx6, a mouse T-Box gene implicated in paraxial mesoderm formation at gastrulation. Dev Biol 1996; 180: 534– 542.

4 Chalamalasetty RB, Garriock RJ, Dunty WCJ, Kennedy MW, Jailwala P, Si H et al. Mesogenin 1 is a master regulator of paraxial presomitic mesoderm differentiation. Development 2014; 141: 4285–4297.

5 Wittler L, Shin E, Grote P, Kispert A, Beckers A, Gossler A et al. Expression of Msgn1 in the presomitic mesoderm is controlled by synergism of WNT signalling and Tbx6. EMBO Rep 2007; 8: 784–789.

6 Hubaud A, Pourquié O. Signalling dynamics in vertebrate segmentation. Nat Rev Mol Cell Biol 2014; 15: 709–721.

7 Cooke J, Zeeman EC. A clock and wavefront model for control of the number of repeated structures during animal morphogenesis. J Theor Biol 1976; 58: 455–476.

8 Pourquié O. Vertebrate somitogenesis. Annu Rev Cell Dev Biol 2001; 17: 311–350.

9 Aulehla A, Herrmann BG. Segmentation in vertebrates: clock and gradient finally joined. Genes Dev 2004; 18: 2060–2067.

10 Oginuma M, Niwa Y, Chapman DL, Saga Y. Mesp2 and Tbx6 cooperatively create periodic patterns coupled with the clock machinery during mouse somitogenesis. Development 2008; 135: 2555 LP – 2562.

11 Moreno TA, Kintner C. Regulation of segmental patterning by retinoic acid signaling during Xenopus somitogenesis. Dev Cell 2004; 6: 205–218.

12 Dubrulle J, McGrew MJ, Pourquié O. FGF signaling controls somite boundary position and regulates segmentation clock control of spatiotemporal Hox gene activation. Cell 2001; 106: 219–232.

13 Christ B, Huang R, Scaal M. Amniote somite derivatives. Dev Dyn 2007; 236: 2382– 2396.

14 Yusuf F, Brand-Saberi B. The eventful somite: patterning, fate determination and cell division in the somite. Anat Embryol (Berl) 2006; 211 Suppl: 21–30.

15 Ibarra-Soria X, Jawaid W, Pijuan-Sala B, Ladopoulos V, Scialdone A, Jörg DJ et al. Defining murine organogenesis at single-cell resolution reveals a role for the leukotriene pathway in regulating blood progenitor formation. Nat Cell Biol 2018; 20: 127–134.

16 Szklarczyk D, Gable AL, Lyon D, Junge A, Wyder S, Huerta-Cepas J et al. STRING v11: Protein-protein association networks with increased coverage, supporting functional discovery in genome-wide experimental datasets. Nucleic Acids Res 2019; 47. doi:10.1093/nar/gky1131.

17 Shannon P, Markiel A, Ozier O, Baliga NS, Wang JT, Ramage D et al. Cytoscape: A software Environment for integrated models of biomolecular interaction networks. Genome Res 2003; 13. doi:10.1101/gr.1239303.

18 Doncheva NT, Morris JH, Gorodkin J, Jensen LJ. Cytoscape StringApp: Network Analysis and Visualization of Proteomics Data. J Proteome Res 2019; 18. doi:10.1021/acs.jproteome.8b00702.

19 Koch F, Scholze M, Wittler L, Schifferl D, Sudheer S, Grote P et al. Antagonistic Activities of Sox2 and Brachyury Control the Fate Choice of Neuro-Mesodermal Progenitors. Dev Cell 2017; 42: 514–526.e7.

20 Xi H, Fujiwara W, Gonzalez K, Jan M, Liebscher S, Van Handel B et al. In Vivo Human Somitogenesis Guides Somite Development from hPSCs. Cell Rep 2017; 18: 1573–1585.

21 Cunningham TJ, Kumar S, Yamaguchi TP, Duester G. Wnt8a and Wnt3a cooperate in the axial stem cell niche to promote mammalian body axis extension. Dev Dyn 2015; 244: 797–807.

22 Tsankov AM, Gu H, Akopian V, Ziller MJ, Donaghey J, Amit I et al. Transcription factor binding dynamics during human ES cell differentiation. Nature 2015; 518: 344–349.

23 Wilkinson DG, Bhatt S, Herrmann BG. Expression pattern of the mouse T gene and its role in mesoderm formation. Nature 1990; 343: 657–659.

24 Tani S, Chung U-I, Ohba S, Hojo H. Understanding paraxial mesoderm development and sclerotome specification for skeletal repair. Exp Mol Med 2020; 52: 1166–1177.

25 Tahara N, Kawakami H, Chen KQ, Anderson A, Peterson MY, Gong W et al. Sall4 regulates neuromesodermal progenitors and their descendants during body elongation in mouse embryos. Development (Cambridge) 2019; 146. doi:10.1242/DEV.177659.

26 Webb SE, Miller AL. Calcium signalling during zebrafish embryonic development. BioEssays. 2000; 22. doi:10.1002/(SICI)1521-1878(200002)22:2<113::AID-BIES3>3.0.CO;2-L.

27 Créton R, Speksnijder JE, Jaffe LF. Patterns of free calcium in zebrafish embryos. J Cell Sci 1998; 111. doi:10.1242/jcs.111.12.1613.

28 Sivak JM, Petersen LF, Amaya E. FGF signal interpretation is directed by sprouty and spred proteins during mesoderm formation. Dev Cell 2005; 8. doi:10.1016/j.devcel.2005.02.011.

29 Nutt SL, Dingwell KS, Holt CE, Amaya E. Xenopus sprouty2 inhibits FGF-mediated gastrulation movements but does not affect mesoderm induction and patterning. Genes Dev 2001; 15. doi:10.1101/gad.191301.

30 Slusarski DC, Pelegri F. Calcium signaling in vertebrate embryonic patterning and morphogenesis. Dev Biol. 2007; 307. doi:10.1016/j.ydbio.2007.04.043.

31 Diaz-Cuadros M, Wagner DE, Budjan C, Hubaud A, Tarazona OA, Donelly S et al. In vitro characterization of the human segmentation clock. Nature 2020; 580: 113–118.

32 Matsuda M, Yamanaka Y, Uemura M, Osawa M, Saito MK, Nagahashi A et al. Recapitulating the human segmentation clock with pluripotent stem cells. Nature 2020; 580: 124–129.

33 Chu L-F, Mamott D, Ni Z, Bacher R, Liu C, Swanson S et al. An In Vitro Human Segmentation Clock Model Derived from Embryonic Stem Cells. Cell Rep 2019; 28: 2247–2255.e5.

34 van den Brink SC, Alemany A, van Batenburg V, Moris N, Blotenburg M, Vivié J et al. Single-cell and spatial transcriptomics reveal somitogenesis in gastruloids. Nature 2020. doi:10.1038/s41586-020-2024-3.

35 Nowotschin S, Ferrer-Vaquer A, Concepcion D, Papaioannou VE, Hadjantonakis A-K. Interaction of Wnt3a, Msgn1 and Tbx6 in neural versus paraxial mesoderm lineage commitment and paraxial mesoderm differentiation in the mouse embryo. Dev Biol 2012; 367: 1–14.

36 Chalamalasetty RB, Dunty WCJ, Biris KK, Ajima R, Iacovino M, Beisaw A et al. The Wnt3a/β-catenin target gene Mesogenin1 controls the segmentation clock by activating a Notch signalling program. Nat Commun 2011; 2: 390.

37 Hofmann M, Schuster-Gossler K, Watabe-Rudolph M, Aulehla A, Herrmann BG, Gossler A. WNT signaling, in synergy with T/TBX6, controls Notch signaling by regulating Dll1 expression in the presomitic mesoderm of mouse embryos. Genes Dev 2004; 18. doi:10.1101/gad.1248604.

38 White PH, Farkas DR, Chapman DL. Regulation of Tbx6 expression by Notch signaling. Genesis 2005; 42: 61–70.

39 Hardy KM, Yatskievych TA, Konieczka J, Bobbs AS, Antin PB. FGF signalling through RAS/MAPK and PI3K pathways regulates cell movement and gene expression in the chicken primitive streak without affecting E-cadherin expression. BMC Dev Biol 2011; 11. doi:10.1186/1471-213X-11-20.

40 Lunn JS, Fishwick KJ, Halley PA, Storey KG. A spatial and temporal map of FGF/Erk1/2 activity and response repertoires in the early chick embryo. Dev Biol 2007; 302. doi:10.1016/j.ydbio.2006.10.014.

41 Aulehla A, Wehrle C, Brand-Saberi B, Kemler R, Gossler A, Kanzler B et al. Wnt3a plays a major role in the segmentation clock controlling somitogenesis. Dev Cell 2003; 4: 395– 406.

42 Ciruna B, Rossant J. FGF Signaling Regulates Mesoderm Cell Fate Specification and Morphogenetic Movement at the Primitive Streak. Dev Cell 2001; 1: 37–49.

43 Chal J, Pourquié O. Making muscle: skeletal myogenesis <em>in vivo</em> and <em>in vitro</em> Development 2017; 144: 2104 LP – 2122.

44 Yamaguchi TP, Harpal K, Henkemeyer M, Rossant J. Fgfr-1 Is Required for Embryonic Growth and Mesodermal Patterning During Mouse Gastrulation. Genes Dev 1994; 8: 3032–3044.

45 Wahl MB, Deng C, Lewandoski M, Pourquié O. FGF signaling acts upstream of the NOTCH and WNT signaling pathways to control segmentation clock oscillations in mouse somitogenesis. Development 2007; 134: 4033–4041.

46 Mujoo K, Krumenacker JS, Murad F. Nitric oxide-cyclic GMP signaling in stem cell differentiation. Free Radic Biol Med. 2011; 51. doi:10.1016/j.freeradbiomed.2011.09.037.

47 Cazzato D, Assi E, Moscheni C, Brunelli S, de Palma C, Cervia D et al. Nitric oxide drives embryonic myogenesis in chicken through the upregulation of myogenic differentiation factors. Exp Cell Res 2014; 320. doi:10.1016/j.yexcr.2013.11.006.

48 Nie S, Chang C. Regulation of Xenopus gastrulation by ErbB signaling. Dev Biol 2007; 303. doi:10.1016/j.ydbio.2006.10.039.

49 Nie S, Chang C. PI3K and Erk MAPK mediate ErbB signaling in Xenopus gastrulation. Mech Dev 2007; 124. doi:10.1016/j.mod.2007.07.005.

50 Pritykin Y, Ghersi D, Singh M. Genome-Wide Detection and Analysis of Multifunctional Genes. PLoS Comput Biol 2015; 11. doi:10.1371/journal.pcbi.1004467.

51 Vilchez D, Boyer L, Lutz M, Merkwirth C, Morantte I, Tse C et al. FOXO4 is necessary for neural differentiation of human embryonic stem cells. Aging Cell 2013; 12. doi:10.1111/acel.12067.

52 Dunwoodie SL, Clements M, Sparrow DB, Sa X, Conlon RA, Beddington RSP. Axial skeletal defects caused by mutation in the spondylocostal dysplasia/pudgy gene Dll3 are associated with disruption of the segmentation clock within the presomitic mesoderm. Development. 2002; 129. doi:10.1242/dev.129.7.1795.

53 Sutherland MJ, Wang S, Quinn ME, Haaning A, Ware SM. Zic3 is required in the migrating primitive streak for node morphogenesis and left-right patterning. Hum Mol Genet 2013; 22. doi:10.1093/hmg/ddt001.

54 Inoue T, Ota M, Mikoshiba K, Aruga J. Zic2 and Zic3 synergistically control neurulation and segmentation of paraxial mesoderm in mouse embryo. Dev Biol 2007; 306. doi:10.1016/j.ydbio.2007.04.003.

55 Ticho BS, Stainier DYR, Fishman MC, Breitbart RE. Three zebrafish MEF2 genes delineate somitic and cardiac muscle development in wild-type and mutant embryos. Mech Dev 1996; 59. doi:10.1016/0925-4773(96)00601-6.

56 Murashima A, Akita H, Okazawa M, Kishigami S, Nakagata N, Nishinakamura R et al. Midline-derived shh regulates mesonephric tubule formation through the paraxial mesoderm. Dev Biol 2014; 386. doi:10.1016/j.ydbio.2013.12.026.

57 Miura S, Davis S, Klingensmith J, Mishina Y. BMP signaling in the epiblast is required for proper recruitment of the prospective paraxial mesoderm and development of the somites. Development 2006; 133. doi:10.1242/dev.02552.

58 Shahbazi MN, Zernicka-Goetz M. Deconstructing and reconstructing the mouse and human early embryo. Nat Cell Biol 2018; 20: 878–887.

59 Yamamoto M, Saijoh Y, Perea-Gomez A, Shawlot W, Behringer RR, Ang S-L et al. Nodal antagonists regulate formation of the anteroposterior axis of the mouse embryo. Nature 2004; 428: 387–392.

60 Arnold SJ, Robertson EJ. Making a commitment: Cell lineage allocation and axis patterning in the early mouse embryo. Nat Rev Mol Cell Biol 2009; 10: 91–103.

61 Rivera-Pérez JA, Magnuson T. Primitive streak formation in mice is preceded by localized activation of Brachyury and Wnt3. Dev Biol 2005; 288: 363–371.

62 Takada S, Stark KL, Shea MJ, Vassileva G, McMahon JA, McMahon AP. Wnt-3a regulates somite and tailbud formation in the mouse embryo. Genes Dev 1994; 8. doi:10.1101/gad.8.2.174.

63 Sun X, Meyers EN, Lewandoski M, Martin GR. Targeted disruption of Fgf8 causes failure of cell migration in the gastrulating mouse embryo. Genes Dev 1999; 13: 1834–1846.

64 Crossley PH, Martin GR. The mouse Fgf8 gene encodes a family of polypeptides and is expressed in regions that direct outgrowth and patterning in the developing embryo. Development 1995; 121. doi:10.1242/dev.121.2.439.

65 Maruoka Y, Ohbayashi N, Hoshikawa M, Itoh N, Hogan BL m., Furuta Y. Comparison of the expression of three highly related genes, Fgf8, Fgf17 and Fgf18, in the mouse embryo. Mech Dev 1998; 74. doi:10.1016/S0925-4773(98)00061-6.

66 Niswander L, Martin GR. Fgf-4 expression during gastrulation, myogenesis, limb and tooth development in the mouse. Development 1992; 114. doi:10.1242/dev.114.3.755.

67 Yang X, Dormann D, Münsterberg AE, Weijer CJ. Cell movement patterns during gastrulation in the chick are controlled by positive and negative chemotaxis mediated by FGF4 and FGF8. Dev Cell 2002; 3. doi:10.1016/S1534-5807(02)00256-3.

68 Boulet AM, Capecchi MR. Signaling by FGF4 and FGF8 is required for axial elongation of the mouse embryo. Dev Biol 2012; 371: 235–245.

69 Diez del Corral R, Storey KG. Opposing FGF and retinoid pathways: a signalling switch that controls differentiation and patterning onset in the extending vertebrate body axis. Bioessays 2004; 26: 857–869.

70 Kudoh T, Wilson SW, Dawid IB. Distinct roles for Fgf, Wnt and retinoic acid in posteriorizing the neural ectoderm. Development 2002; 129: 4335–4346.

71 Yamaguchi TP, Takada S, Yoshikawa Y, Wu N, McMahon AP. T (Brachyury) is a direct target of Wnt3a during paraxial mesoderm specification. Genes Dev 1999; 13: 3185–3190.

72 Wittler L, Shin EH, Grote P, Kispert A, Beckers A, Gossler A et al. Expression of Msgn1 in the presomitic mesoderm is controlled by synergism of WNT signalling and Tbx6. EMBO Rep 2007; 8: 784–789.

73 Sudheer S, Liu J, Marks M, Koch F, Anurin A, Scholze M et al. Different Concentrations of FGF Ligands, FGF2 or FGF8 Determine Distinct States of WNT-Induced Presomitic Mesoderm. Stem Cells 2016; 34. doi:10.1002/stem.2371.

74 Martin BL, Kimelman D. Brachyury establishes the embryonic mesodermal progenitor niche. Genes Dev 2010; 24: 2778–2783.

75 Aulehla A, Pourquié O. Signaling gradients during paraxial mesoderm development. Cold Spring Harb Perspect Biol 2010; 2: a000869.

76 Aulehla A, Wiegraebe W, Baubet V, Wahl MB, Deng C, Taketo M et al. A beta-catenin gradient links the clock and wavefront systems in mouse embryo segmentation. Nat Cell Biol 2008; 10: 186–193.

77 Dale JK, Malapert P, Chal J, Vilhais-Neto G, Maroto M, Johnson T et al. Oscillations of the snail genes in the presomitic mesoderm coordinate segmental patterning and morphogenesis in vertebrate somitogenesis. Dev Cell 2006; 10: 355–366.

78 Rifes P, Carvalho L, Lopes C, Andrade RP, Rodrigues G, Palmeirim I et al. Redefining the role of ectoderm in somitogenesis: a player in the formation of the fibronectin matrix of presomitic mesoderm. Development 2007; 134: 3155–3165.

79 Yabe T, Takada S. Molecular mechanism for cyclic generation of somites: Lessons from mice and zebrafish. Dev Growth Differ 2016; 58: 31–42.

80 Bessho Y, Hirata H, Masamizu Y, Kageyama R. Periodic repression by the bHLH factor Hes7 is an essential mechanism for the somite segmentation clock. Genes Dev 2003; 17: 1451–1456.

81 Matsumiya M, Tomita T, Yoshioka-Kobayashi K, Isomura A, Kageyama R. ES cell-derived presomitic mesoderm-like tissues for analysis of synchronized oscillations in the segmentation clock. Development 2018; 145. doi:10.1242/dev.156836.

82 Niwa H, Ogawa K, Shimosato D, Adachi K. A parallel circuit of LIF signalling pathways maintains pluripotency of mouse ES cells. Nature 2009; 460: 118–122.

83 Harima Y, Kageyama R. Oscillatory links of Fgf signaling and Hes7 in the segmentation clock. Curr Opin Genet Dev 2013; 23: 484–490.

84 Oginuma M, Takahashi Y, Kitajima S, Kiso M, Kanno J, Kimura A et al. The oscillation of Notch activation, but not its boundary, is required for somite border formation and rostral-caudal patterning within a somite. Development 2010; 137: 1515–1522.

85 Saga Y. Segmental border is defined by the key transcription factor Mesp2, by means of the suppression of Notch activity. Dev Dyn 2007; 236: 1450–1455.

86 Sasaki N, Kiso M, Kitagawa M, Saga Y. The repression of Notch signaling occurs via the destabilization of mastermind-like 1 by Mesp2 and is essential for somitogenesis. Development 2011; 138: 55–64.

87 Yasuhiko Y, Haraguchi S, Kitajima S, Takahashi Y, Kanno J, Saga Y. Tbx6-mediated Notch signaling controls somite-specific Mesp2 expression. Proc Natl Acad Sci U S A 2006; 103: 3651–3656.

88 Takahashi Y, Takagi A, Hiraoka S, Koseki H, Kanno J, Rawls A et al. Transcription factors Mesp2 and Paraxis have critical roles in axial musculoskeletal formation. Developmental Dynamics 2007; 236. doi:10.1002/dvdy.21178.

89 Tajbakhsh S, Borello U, Vivarelli E, Kelly R, Papkoff J, Duprez D et al. Differential activation of Myf5 and MyoD by different Wnts in explants of mouse paraxial mesoderm and the later activation of myogenesis in the absence of Myf5. Development 1998; 125: 4155–4162.

90 Ikeya M, Takada S. Wnt signaling from the dorsal neural tube is required for the formation of the medial dermomyotome. Development 1998; 125: 4969–4976.

91 Wallingford JB, Fraser SE, Harland RM. Convergent extension: The molecular control of polarized cell movement during embryonic development. Dev Cell. 2002; 2. doi:10.1016/S1534-5807(02)00197-1.

92 Schlessinger J. SH2/SH3 signaling proteins. Curr Opin Genet Dev 1994; 4. doi:10.1016/0959-437X(94)90087-6.

93 Dorey K, Amaya E. FGF signalling: diverse roles during early vertebrate embryogenesis. Development 2010; 137: 3731–3742.

94 Lee H-J, Göring W, Ochs M, Mühlfeld C, Steding G, Paprotta I et al. Sox15 Is Required for Skeletal Muscle Regeneration. Mol Cell Biol 2004; 24. doi:10.1128/mcb.24.19.8428-8436.2004.

95 Meeson AP, Shi X, Alexander MS, Williams RS, Allen RE, Jiang N et al. Sox15 and Fhl3 transcriptionally coactivate Foxk1 and regulate myogenic progenitor cells. EMBO Journal 2007; 26. doi:10.1038/sj.emboj.7601635.

96 van Nes J, de Graaff W, Lebrin F, Gerhard M, Beck F, Deschamps J. The Cdx4 mutation affects axial development and reveals an essential role of Cdx genes in the ontogenesis of the placental labyrinth in mice. Development 2006; 133. doi:10.1242/dev.02216.

97 Pan H, Gustafsson MK, Aruga J, Tiedken JJ, Jennifer JC, Emerson CP. A role for Zic1 and Zic2 in Myf5 regulation and somite myogenesis. Dev Biol 2011; 351. doi:10.1016/j.ydbio.2010.12.037.

98 Odaka YS, Tohmonda T, Toyoda A, Aruga J. An Evolutionarily Conserved Mesodermal Enhancer in Vertebrate Zic3. Sci Rep 2018; 8. doi:10.1038/s41598-018-33235-y.

99 Reshef R, Maroto M, Lassar AB. Regulation of dorsal somitic cell fates: BMPs and Noggin control the timing and pattern of myogenic regulator expression. Genes Dev 1998; 12: 290–303.

100 Kondo N, Tohnai G, Sahashi K, Iida M, Kataoka M, Nakatsuji H et al. DNA methylation inhibitor attenuates polyglutamine-induced neurodegeneration by regulating Hes5. EMBO Mol Med 2019; 11. doi:10.15252/emmm.201708547.

101 Long F, Shi H, Li P, Guo S, Ma Y, Wei S et al. A SMOC2 variant inhibits BMP signaling by competitively binding to BMPR1B and causes growth plate defects. Bone 2021; 142. doi:10.1016/j.bone.2020.115686.

102 Gomez-Puerto MC, Iyengar PV, García de Vinuesa A, ten Dijke P, Sanchez-Duffhues G. Bone morphogenetic protein receptor signal transduction in human disease. Journal of Pathology. 2019; 247. doi:10.1002/path.5170.

103 Ma X, Su P, Yin C, Lin X, Wang X, Gao Y et al. The roles of foxo transcription factors in regulation of bone cells function. Int J Mol Sci. 2020; 21. doi:10.3390/ijms21030692.

104 Sanchez AMJ, Candau RB, Bernardi H. FoxO transcription factors: Their roles in the maintenance of skeletal muscle homeostasis. Cellular and Molecular Life Sciences. 2014; 71. doi:10.1007/s00018-013-1513-z.

105 Teixeira CC, Liu Y, Thant LM, Pang J, Palmer G, Alikhani M. Foxo1, a novel regulator of osteoblast differentiation and skeletogenesis. Journal of Biological Chemistry 2010; 285. doi:10.1074/jbc.M109.079962.

106 Yu S, Guo J, Sun Z, Lin C, Tao H, Zhang Q et al. BMP2-dependent gene regulatory network analysis reveals Klf4 as a novel transcription factor of osteoblast differentiation. Cell Death Dis 2021; 12. doi:10.1038/s41419-021-03480-7.

107 Lyu Y, Nakano K, Davis RR, Tepper CG, Campbell M, Izumiya Y. ZIC2 Is Essential for Maintenance of Latency and Is a Target of an Immediate Early Protein during Kaposi’s Sarcoma-Associated Herpesvirus Lytic Reactivation. J Virol 2017; 91. doi:10.1128/jvi.00980-17.

108 Salhotra A, Shah HN, Levi B, Longaker MT. Mechanisms of bone development and repair. Nat Rev Mol Cell Biol. 2020; 21. doi:10.1038/s41580-020-00279-w.

109 Juneja SC, Vonica A, Zeiss C, Lezon-Geyda K, Yatsula B, Sell DR et al. Deletion of mecom in mouse results in early-onset spinal deformity and osteopenia. Bone 2014; 60. doi:10.1016/j.bone.2013.11.020.

110 Dey BK, Mueller AC, Dutta A. Long non-coding rnas as emerging regulators of differentiation, development, and disease. Transcription. 2014; 5. doi:10.4161/21541272.2014.944014.

111 Piasecka A, Sekrecki M, Szcześniak MW, Sobczak K. MEF2C shapes the microtranscriptome during differentiation of skeletal muscles. Sci Rep 2021; 11. doi:10.1038/s41598-021-82706-2.

112 Chen X, Gao B, Ponnusamy M, Lin Z, Liu J. MEF2 signaling and human diseases. Oncotarget. 2017; 8. doi:10.18632/oncotarget.22899.

113 Arnold MA, Kim Y, Czubryt MP, Phan D, McAnally J, Qi X et al. MEF2C Transcription Factor Controls Chondrocyte Hypertrophy and Bone Development. Dev Cell 2007; 12. doi:10.1016/j.devcel.2007.02.004.

114 Kan C, Chen L, Hu Y, Ding N, Li Y, McGuire TL et al. Gli1-labeled adult mesenchymal stem/progenitor cells and hedgehog signaling contribute to endochondral heterotopic ossification. Bone 2018; 109. doi:10.1016/j.bone.2017.06.014.

115 Park HL, Bai C, Platt KA, Matise MP, Beeghly A, Hui CC et al. Mouse Gli1 mutants are viable but have defects in SHH signaling in combination with a Gli2 mutation. Development 2000; 127. doi:10.1242/dev.127.8.1593.

116 Houtmeyers R, Souopgui J, Tejpar S, Arkell R. The ZIC gene family encodes multi-functional proteins essential for patterning and morphogenesis. Cellular and Molecular Life Sciences. 2013; 70. doi:10.1007/s00018-013-1285-5.

117 Rocca-Serra P, Brazma A, Parkinson H, Sarkans U, Shojatalab M, Contrino S et al. ArrayExpress: A public database of gene expression data at EBI. C R Biol 2003; 326. doi:10.1016/j.crvi.2003.09.026.

118 Sarkans U, Füllgrabe A, Ali A, Athar A, Behrangi E, Diaz N et al. From ArrayExpress to BioStudies. Nucleic Acids Res 2021; 49. doi:10.1093/nar/gkaa1062.

119 Barrett T, Wilhite SE, Ledoux P, Evangelista C, Kim IF, Tomashevsky M et al. NCBI GEO: Archive for functional genomics data sets - Update. Nucleic Acids Res 2013; 41. doi:10.1093/nar/gks1193.

120 Kim D, Paggi JM, Park C, Bennett C, Salzberg SL. Graph-based genome alignment and genotyping with HISAT2 and HISAT-genotype. Nat Biotechnol 2019; 37. doi:10.1038/s41587-019-0201-4.

121 Trapnell C, Williams BA, Pertea G, Mortazavi A, Kwan G, Van Baren MJ et al. Transcript assembly and quantification by RNA-Seq reveals unannotated transcripts and isoform switching during cell differentiation. Nat Biotechnol 2010; 28: 511–515.

122 Szklarczyk D, Franceschini A, Wyder S, Forslund K, Heller D, Huerta-Cepas J et al. STRING v10: Protein-protein interaction networks, integrated over the tree of life. Nucleic Acids Res 2015; 43. doi:10.1093/nar/gku1003.

123 Dennis G, Sherman BT, Hosack DA, Yang J, Gao W, Lane HC et al. DAVID: Database for Annotation, Visualization, and Integrated Discovery. Genome Biol 2003; 4. doi:10.1186/gb-2003-4-9-r60.

