## Supplementary figures and tables for "Identification of novel evolutionarily conserved genes and pathways in human and mouse musculoskeletal progenitors": Supplementary Figures_29122023.pdf

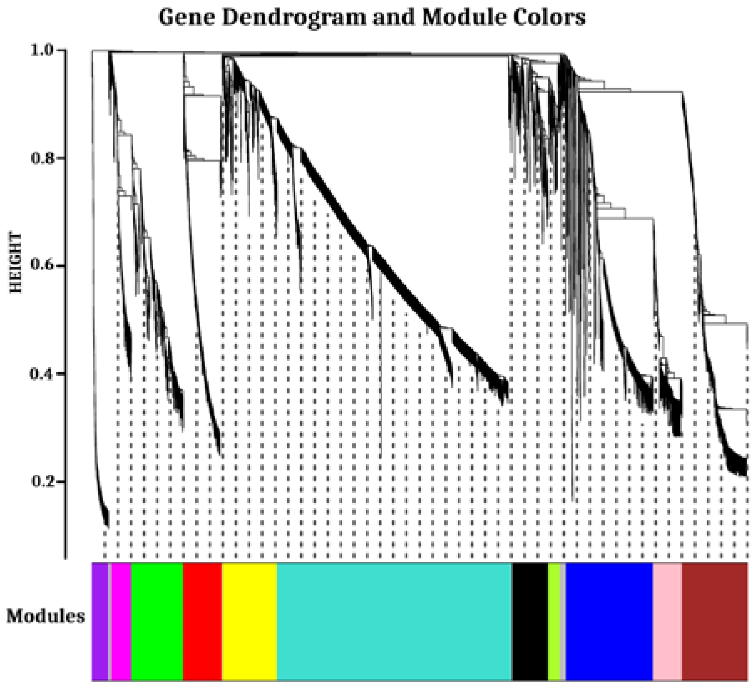


**Figure S1**: WGCNA clustering of DEGs involved in the paraxial mesoderm development of mouse

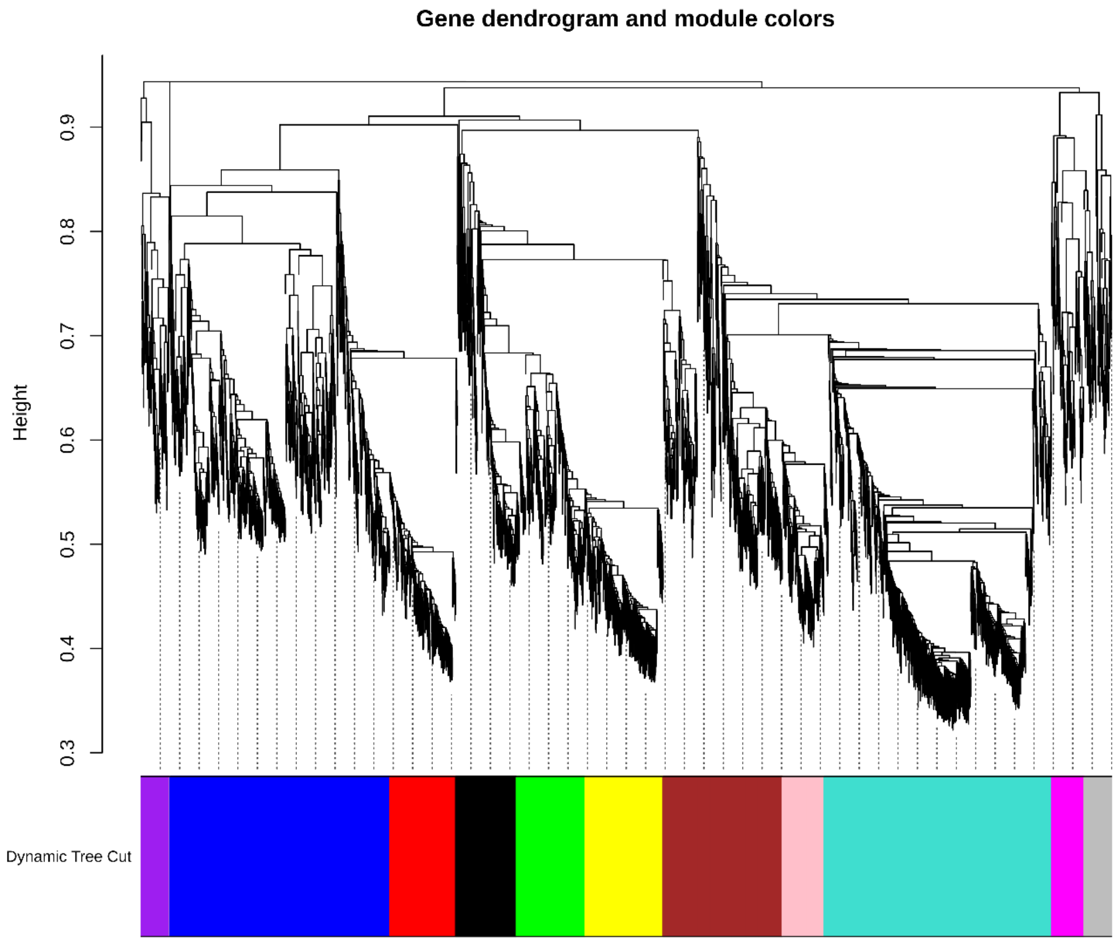


**Figure S2**: WGCNA clustering of DEGs involved in the paraxial mesoderm development of human

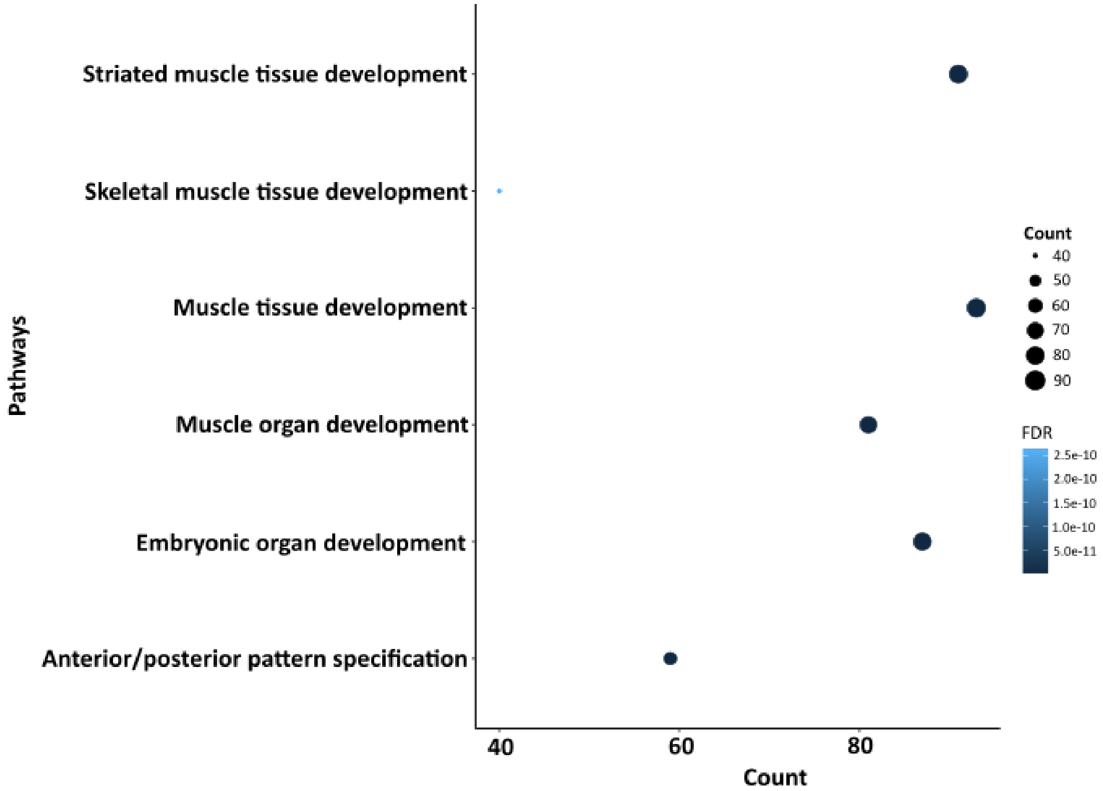


**Figure S3.** Biological process mouse of evolutionarily conserved genes

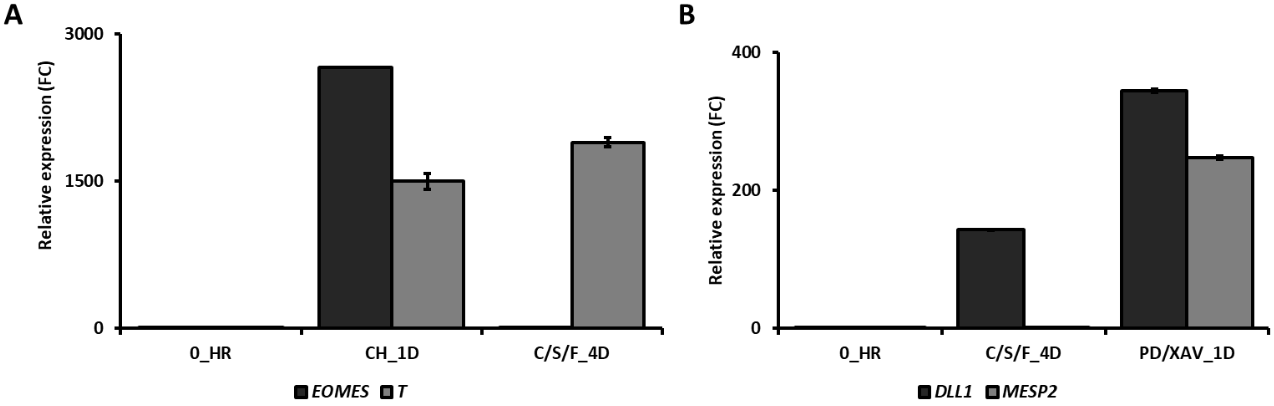


**Figure S4**: qRT-PCR data of PS and somite markers of indicated samples. Data are Mean ± s.d., n=2

| Gene | Primer_F | Primer_R |
|---|---|---|
| *ACTB* | TCAAGATCATTGCTCCTCCTGAG | ACATCTGCTGGAAGGTGGACA |
| *BMP2* | GCAGCTTCCACCATGAAGA | GCAGCTTCCACCATGAAGA |
| *BMP5* | AGGGGATGGACGCAGTATC | TGAAGAAGGCCACCATGAA |
| *BMP7* | CCATCGAGAGTTCCGGTTT | AAGCGTTCCCGGATGTAGT |
| *DLL1* | CTGCAACCAGGACCTGAACT | CCGGCAAGAGCAAGTGTAG |
| *EOMES* | CGGCCTCTGTGGCTCAAA | AAGGAAACATGCGCCTGC |
| *GLI1* | CTCCCCACCAGAGAATGGA | CGAAGTACCACCCCCTCTG |
| *HES5* | AGCGAAGGCTACTCGTGGT | GCCGCTGGAAGTGGTACAG |
| *MEF2C* | GCTGTTCCACCTCCCAACT | TGGCAATAGGTTGGGGTTT |
| *MEOX1* | GATGACTACGGGGTGCTTG | TCCTGGTTGTCTGAACTCTCC |
| *MSGN1* | CTGCACACCCTCCGGAATT | CTCTGCCGCGGTTAAGGAG |
| *NOG* | TCGAACACCCAGACCCTATC | TGAAGCCTGGGTCGTAGTG |
| *RIPPLY1* | GCTCCAGACCTAGCACAAGC | GCCTCCAGAGACAAGTTCCTC |
| *T* | CCTTGCTCACACCTGCAGTAGC | GGCCAACTGCATCATCTCCA |
| *TBX6* | AGCCTGTGTCTTTCCATCGT | GCTGCCCGAACTAGGTGTAT |
| *ZIC3* | GCTGCCAGTTCAGGCTATG | AGGAAGTCCAGGGTTGTGG |

**Table S6:** List of Primers
